# PRMT5 regulates ATF4 transcript splicing and oxidative stress response

**DOI:** 10.1101/2022.02.02.478789

**Authors:** Magdalena M. Szewczyk, Genna M. Luciani, Victoria Vu, Alex Murison, David Dilworth, Mathieu Lupien, Cheryl H Arrowsmith, Mark D. Minden, Dalia Barsyte-Lovejoy

## Abstract

Protein methyltransferase 5 (PRMT5) symmetrically dimethylates arginine residues leading to regulation of transcription and splicing programs. Although PRMT5 has emerged as an attractive oncology target, the molecular determinants of PRMT5 dependency in cancer remain incompletely understood. Our transcriptomic analysis identified PRMT5 regulation of the activating transcription factor 4 (ATF4) pathway in acute myelogenous leukemia (AML). PRMT5 inhibition resulted in the expression of unstable, intron-retaining ATF4 mRNA that is detained in the nucleus. Concurrently, the decrease in the spliced cytoplasmic transcript of ATF4 led to lower levels of ATF4 protein and downregulation of ATF4 target genes. Upon loss of functional PRMT5, cells with low ATF4 displayed increased oxidative stress, growth arrest, and cellular senescence. Interestingly, leukemia cells with EVI1 oncogene overexpression demonstrated dependence on PRMT5 function. EVI1 and ATF4 regulated gene signatures were inversely correlated. We show that EVI1-high AML cells have reduced ATF4 levels, elevated baseline reactive oxygen species and increased sensitivity to PRMT5 inhibition. Thus, EVI1-high cells demonstrate dependence on PRMT5 function and regulation of oxidative stress response. Overall, our findings identify the PRMT5-ATF4 axis to be safeguarding the cellular redox balance that is especially important in high oxidative stress states, such as those that occur with EVI1 overexpression.

## Introduction

Protein arginine methyltransferases (PRMT) mono or dimethylate arginines in an asymmetric or symmetric manner and regulate various biological processes, such as transcription, splicing, translation, and cell signaling (1). The PRMT5 enzyme is responsible for most of the cellular symmetric dimethylation of arginines (SDMA) (2, 3), of histones (4), transcription factors (5), signaling proteins (6), spliceosome assembly factors (7), as well as numerous RNA binding and processing-associated proteins (8-10). This diversity of PRMT5 substrates explains the broad range of PRMT5 functions that include the regulation of DNA damage repair, splicing, transcription, translation, metabolism, and stress response (11, 12).

The regulation of splicing by PRMT5 has been the focus of several recent investigations. Initial observations highlighted PRMT5 control of MDM4 exon skipping and levels of P53 (7); PRMT5 inhibition leads to activation of the P53 pathway and subsequent cell death (13). In addition, splice isoform switch of the histone acetyltransferase KAT5 mediated by PRMT5 is important in regulating the DNA damage response and cell death (14). Finally, methylation of a key splicing factor SRSF1 by PRMT5 affects SRSF1 binding to splice sites resulting in broad changes in splicing programs revealing pleiotropic, cell-type, and state-specific regulatory responses regulated by PRMT5 (10).

PRMT5 is overexpressed in many cancer types, including lymphoma, glioma, breast, and lung cancer (15, 16). It has been reported to be essential for normal hematopoiesis (17, 18) as well as the maintenance of leukemia and lymphoma (15). For a subset of acute myeloid leukemia (AML) with MLL-rearrangement, PRMT5 inhibition results in defective differentiation (19). AML and myelodysplasia patient cells frequently have mutations in splicing factors that confers synthetic lethality upon PRMT5 inhibition (8). Overall, the existing data indicate a critical role for PRMT5 in regulation of the splicing program, which is often dysregulated in cancer. As such, PRMT5 is emerging as an attractive therapeutic target with active tool compound development (20, 21) and ongoing, early phase clinical studies (1, 22).

To better understand the mechanistic role of PRMT5, we investigated downstream transcriptome regulation by PRMT5, focusing on poor prognosis forms of AML that are dependent on PRMT5 function. One such subclass of AML has a rearrangement of chromosome 3q21 and is dependent on ecotropic virus integration site 1 (EVI1) factor-driven gene expression programs affecting stemness, apoptotic response, and differentiation (23-25). In this group of AML, our transcriptomic analysis identified activating transcription factor 4 (ATF4) as a PRMT5 target. ATF4 controls gene expression associated with metabolic and redox processes and plays a major role in the unfolded protein response (UPR) (26, 27). PRMT5 inhibition resulted in the expression of a highly unstable ATF4 splice variant mRNA that was detained in the nucleus. Consequently, ATF4 protein levels were downregulated, as was the downstream transcriptional ATF4 program associated with the oxidative stress response. To understand the significance of PRMT5-ATF4 regulation in the context of EVI1, we determined that EVI1 overexpression also increased levels of reactive oxygen species, and EVI1-high in leukemia cells had reduced ATF4 levels and increased oxidative stress, subsequently sensitizing these cells to PRMT5 inhibition.

## Results

### The catalytic activity of PRMT5 is required for EVI1 AML proliferation

Various leukemia genetic backgrounds display differing dependence on PRMT5. We have previously shown that SRSF2 and SF3B1 splicing factor mutations preferentially sensitize cells to PRMT inhibition (8) and subsequently identified additional patient subgroups with dependency on PRMT5 catalytic function. One of these groups were leukemia cells with the recurrent cytogenetic abnormality inversion of chromosome 3 (inv3 and t(3;3)), that results in the constitutive expression of the proto-oncogene EVI1. Similar to SRSF2 mutant leukemia, proliferation of primary patient cells with inv(3) and a derived line OCI-AML-20 (28) were inhibited by PRMT5 inhibition (Fig. 1a-b, Suppl. Fig. 1a-c). Among the primary samples tested, the P53 mutant inv(3) sample 80309 was the least growth inhibited (Fig. 1a, Suppl Fig. 1c). Given that there was only a minimal induction of apoptosis in EVI1-high cells (Suppl Fig. 1d), we further investigated potential mechanisms underlying cell proliferation suppression by PRMT5 catalytic function inhibition. Treatment with two chemically distinct PRMT5 inhibitors (PRMT5i), GSK591 (21) and LLY283 (20) but not the negative control compound SGC2096, resulted in a decreased proportion of S-phase cells (Fig. 1c, Suppl. Fig. 1e-f) and increased cell quiescence (Fig. 1d-e). We also investigated if the proliferation defect is due to induction of senescence and found that exposure to PRMT5 inhibitor for ten days significantly increased levels of β-galactosidase activity in OCI-AML-20 cells (Fig. 1f). These results indicate that PRMT5 catalytic activity is required for EVI1 AML cell cycle progression and PRMT5 inhibition leads to cell cycle arrest and senescence.

**Fig. 1.**
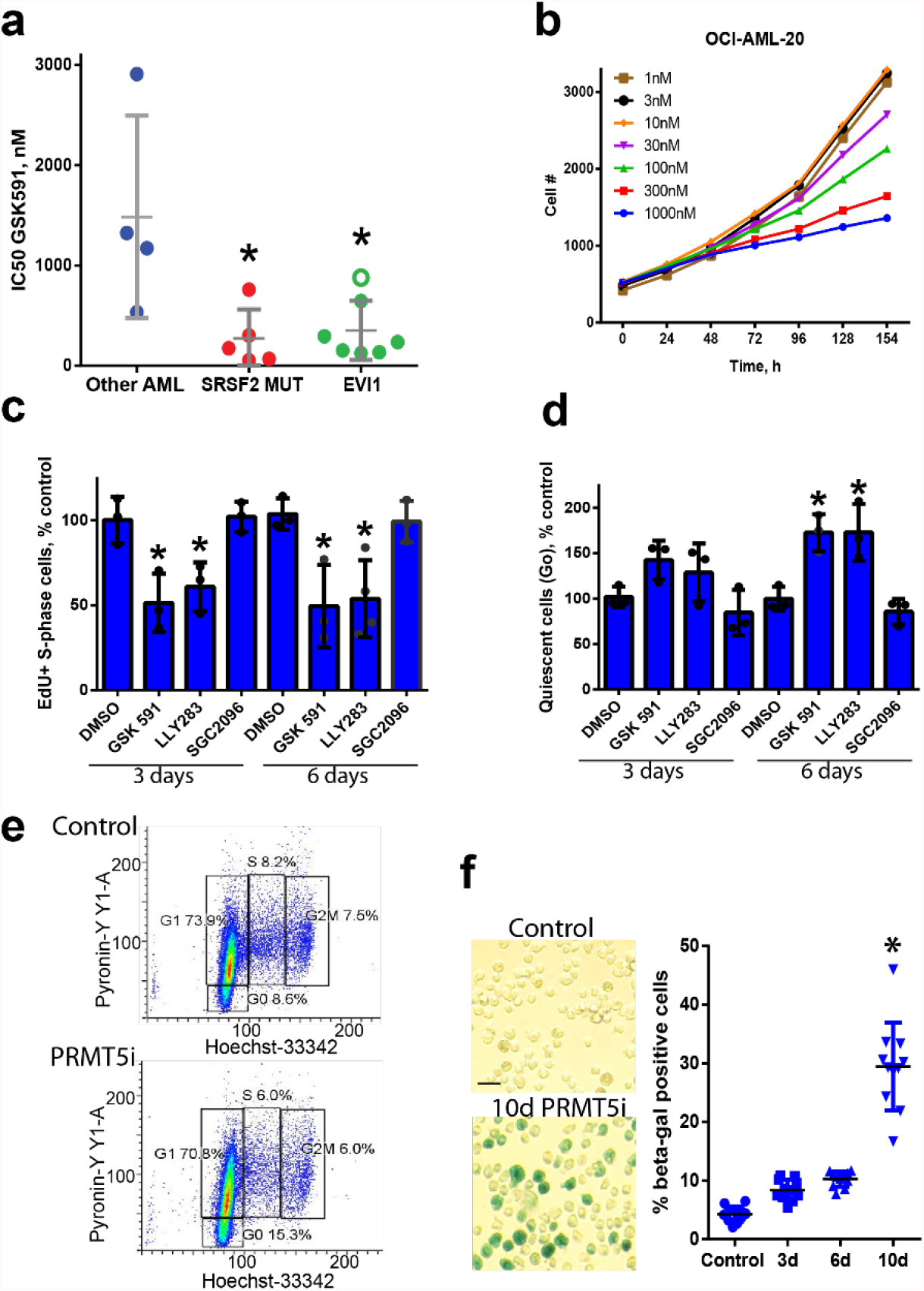
PRMT5 inhibition leads to a deficiency in cell proliferation, S-phase decrease, increased quiescence, and cell senescence. **a)** Leukemia cell growth impairment by PRMT5 inhibition. IC50 values for PRMT5 inhibitor GSK591 in EVI1-high, SRSF2 mutant, and other genetic background leukemia patient samples (Other AML). Star indicates p values for SFRSF2mut of 0.016 and p=0.007 for EVI1-high cells excluding the mutant TP53 sample (open green circle) (Kruskal-Wallis, posthoc Dunn’s test). **b)** Kinetics of cell growth of EVI1-high OCI-AML-20 cells in response to GSK591 (means of 3 replicates shown). **c)** PRMT5 inhibition results in decreased S-phase frequency (1 μM, GSK591, LLY283, and SGC2096 negative control compound). N=3, data represented as mean±SD, * p<0.05 relative to time-matched control. **d)** Increased quiescence in EVI-high OCI-AML-20 treated with PRMT5 inhibitors as in C. N=3, data represented as mean±SD, * p<0.05 relative to time-matched control. **e)** Flow cytometry plots illustrating G0 quiescent populations for data in d). **f)** PRMT5 inhibition (GSK591 1 μM) induces senescence in OCI-AML-20 cells as shown by β-galactosidase staining and quantitation on the right (N=3, all individual data points shown, * p<0.05, mean±SD).

### PRMT5 regulates ATF4 transcriptional program

To determine the underlying mechanism of PRMT5i response in EVI1 rearranged leukemia, RNAseq was performed for PRMT5i GSK598 treated and control OCI-AML-20 cells. Changes in gene expression at the 6-day timepoint were investigated to capture the program associated with the senescence response (Fig. 1f). While overall, there was a greater number of upregulated than downregulated transcripts (Fig. 2a), significantly enriched pathways were evident only in the downregulated gene set. The pathways associated with unfolded protein response (UPR) constituted five out of seven of the most significantly enriched pathways in the downregulated gene set (Fig. 2b). Further analysis using Transcriptional Regulatory Relationships Unraveled by Sentence-based Text-mining (TRRUST), an extensive database of transcription factors and their targets (29), identified ATF4, which is a critical transcription factor of the UPR pathway as being most significantly represented regulator of the identified PRMT5i downmodulated gene set (Fig. 2c). Validation of differentially expressed gene findings by qRT-PCR confirmed that PRMT5 inhibition resulted in lower mRNA expression levels of ATF4 and its target genes ASNS, PSAT, TRIB3, CHOP at six days and also at the 3-day time point (Fig. 2d). PHGDH, SHMT2, amino acid, and metabolite transporters such as SLC7A5 and SLC7A1 were downregulated at six days (Fig. 2d). Several transcripts were validated as being upregulated upon PRMT5 inhibition (Suppl. Fig. 2a). Other regulators of non-ATF4 branches of the UPR, such as activating transcription factor 6 (ATF6), PERK, or spliced X-box binding protein 1 (XBP1), were not affected by PRMT5 inhibition. (Suppl. Fig. 2b). These results suggest that PRMT5 activity is required for maintaining the levels of ATF4 and its downstream-regulated programs.

**Fig. 2.**
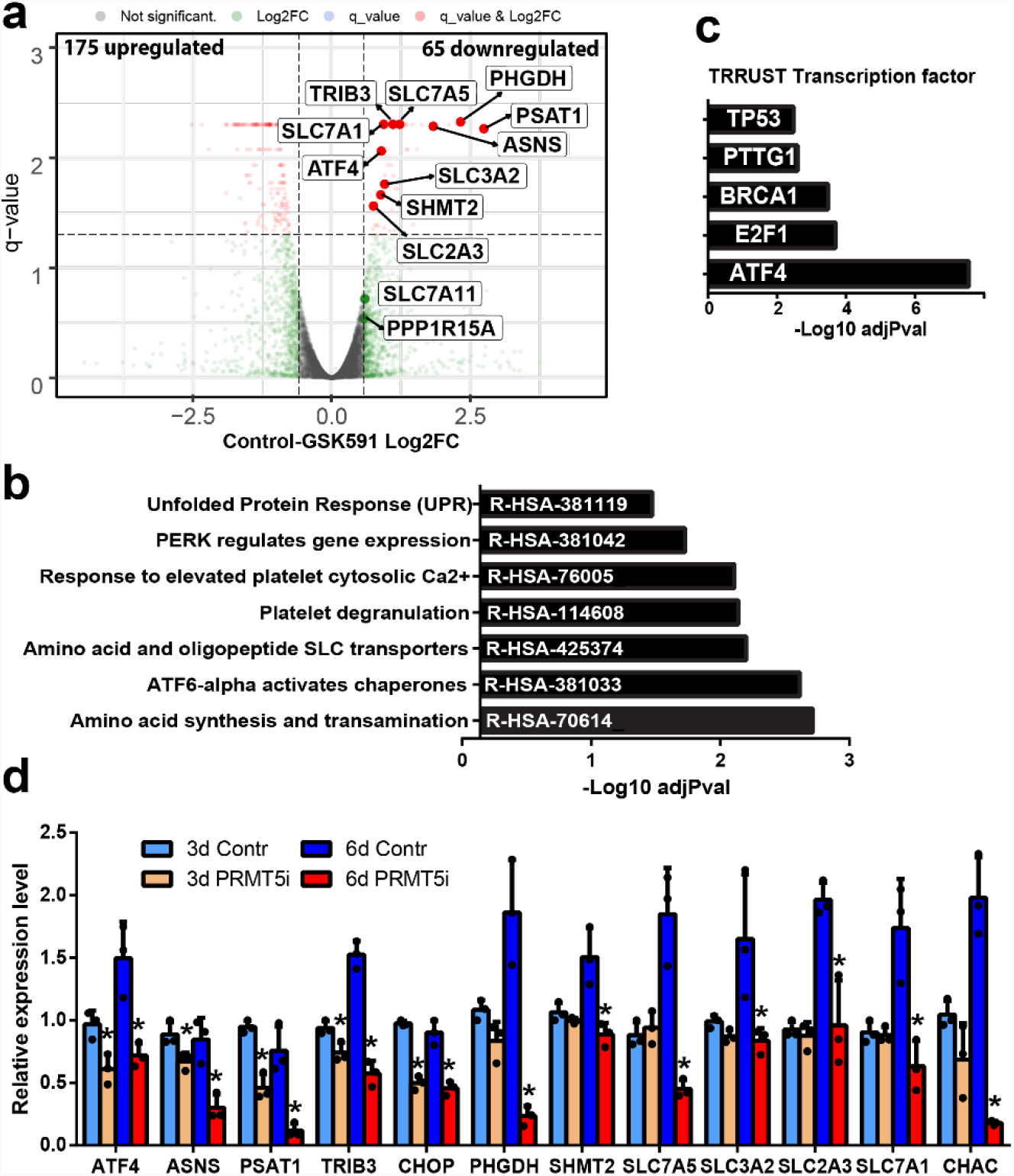
PRMT5 regulates gene expression and ATF4 transcriptional program. **a)** Transcripts upregulated and downregulated by PRMT5i in OCI-AML-20 as illustrated by RNAseq volcano plots for DMSO control and GSK591 (6 days, 1 μM GSK591). **b)** Reactome pathways significantly enriched in the PRMT5i downregulated transcript set. **c)** Significantly enriched transcription factor binding sites in the downregulated gene set as identified by TRRUST (transcriptional regulatory relationships unraveled by sentence-based text-mining) analysis. **d)** Validation of significantly downregulated transcripts, 3 and 6 days of PRMT5 inhibition. N=3, mean±SD shown, * p<0.05 of the day-matched control sample. Validation was performed in OCI-AML-20 cells using PRMT5 inhibitor LLY283 at 100 nM.

### PRMT5 differentially regulates ATF4 splice variants

Previous studies have implicated PRMT5 in splicing regulation (7, 8, 30, 31). In keeping with this, we observed that PRMT5 inhibition led to intron retention (598 events) and exon skipping (1394 events). Genes affected by these alternative splicing events were mainly associated with RNA regulatory processes. (Fig. 3a, Suppl. Fig. 3a). Since ATF4 mRNA levels were reduced by PRMT5 inhibition, we asked if there was any difference in the ATF4 mRNA splice forms. ATF4 has two splice variants: variant 1 (V1, NM_001675.4) has a retained first intron between nucleic acids from the transcription start site to the translation start site (1-887), while variant 2 that encodes for canonical ATF4 (V2, NM_182810.3) has the first intron spliced out and a shorter 5’ end of 282 nucleosides (Fig. 3b). Following treatment with PRMT5i, there was an increase in the level of V1 and concurrent reduction in the level of V2 mRNA at six days (Fig. 3c). This was observed in OCI-AML-20 and another well-utilized EVI1 AML cell line with the same genetic abnormality of inv3, UCSD-AML-1 (Fig. 3c). Inspection of *ATF4* gene exon-intron structure identified a weak splice donor site (Fig. 3b) that varied from the consensus splice site (32). Such non-consensus, weak splice donor sites have been reported to be dependent on PRMT5 function (7). Evolutionary sequence analysis also indicated that the non-consensus splice site is conserved in vertebrate *ATF4* genes (Fig 3b).

**Fig. 3.**
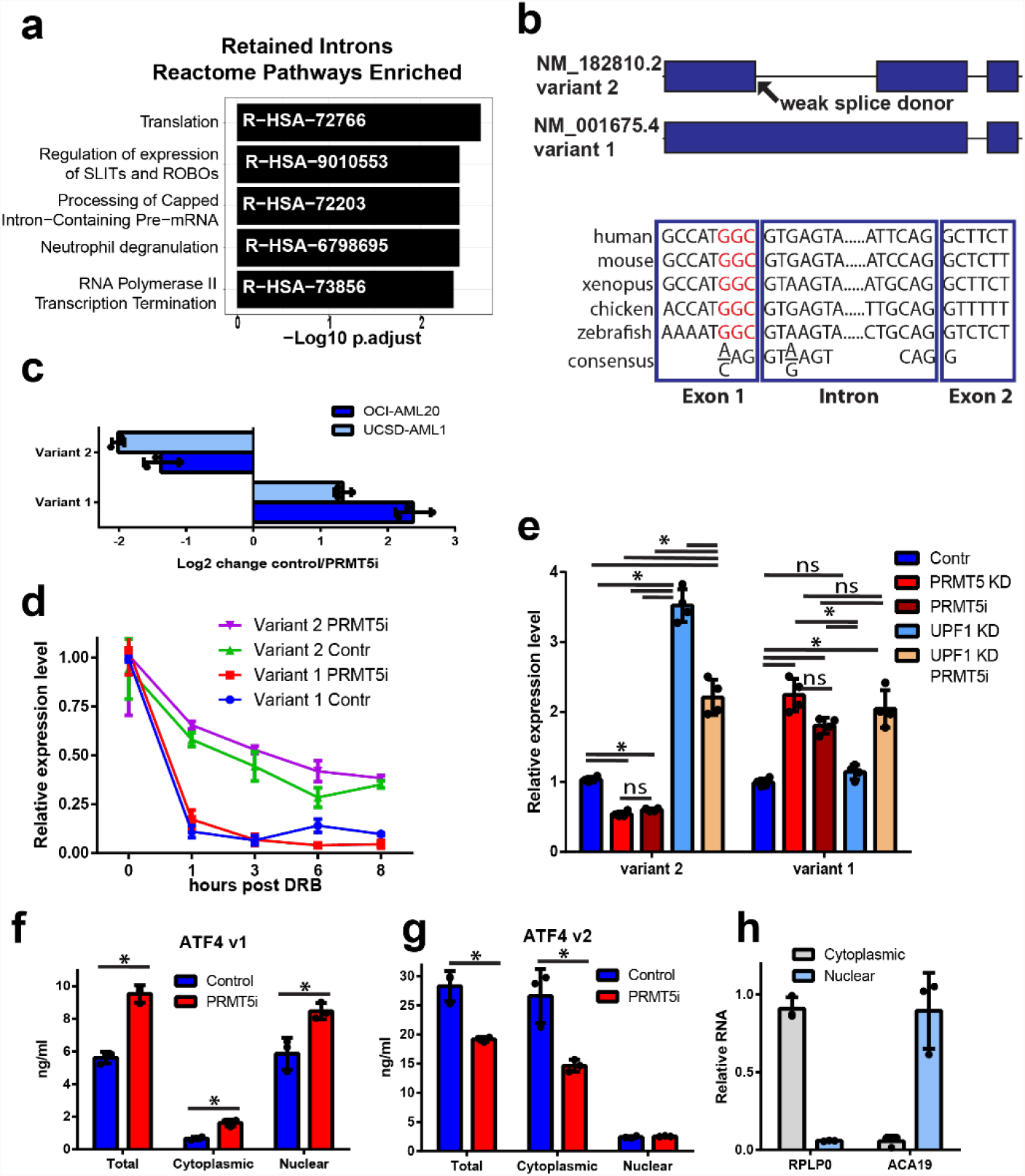
PRMT5 regulates splicing programs and ATF4 splice variant switch. **a)** Significantly enriched Reactome pathways in PRMT5 inhibition regulated intron retention events. **b)** A schematic of *ATF4* gene illustrating the spliced variant 2 and retention of intron 1 that generates variant 1. The evolutionary conservation of weak 5’ splice donor sites in intron 1 of the *ATF4* gene is illustrated below, red non-consensus sequence. **c)** Fold change in *ATF4* variant 1 and 2 transcripts upon PRMT5 inhibition by 100 nM LLY283. Normalized expression fold change is shown where negative values indicate downregulation and positive – upregulation due to PRMT5 inhibition (N=3, mean±SD). **d)** Variant 1 and variant 2 transcripts have differential stability. UCSD-AML-1 cells, pretreated with LLY283 100 nM for 6 days, were exposed to 5,6-dichloro-1-beta-D-ribofuranosylbenzimidazole (DRB) transcription inhibitor, and transcript levels were assessed by PCR. Normalized relative expression level is plotted, N=3, mean±SEM shown. **e)** Inhibition of nonsense-mediated decay (UPF1 knockdown) affected the variant 2 but not variant 1 transcript levels (PRMT5i, LLY283 100 nM, 6 days, UCSD-AML-1 cells). N=3, * p<0.05, meas±SD, two-way ANOVA. **f-g)** Nuclear and cytoplasmic fractions of V1 and V2 of ATF4 as determined by quantitative standard curve-based RT-PCR (N=3, * p<0.05, mean±SD). **h)** Fractionation quality controls of nuclear ACA19 and cytoplasmic RPLP0 RNAs normalized to a nuclear or cytoplasmic fraction.

While PRMT5i treatment was associated with decreased levels of ATF4 V2 mRNA and relative increase of V1 mRNA (Fig. 3c), the levels of intronic transcript appeared to be minor (Suppl. Fig. 3b), thus raising a possibility that V1 transcript is unstable. To assess transcript stability, cells were treated with the transcription inhibitor 5,6-dichloro-1-beta-D-ribofuranosylbenzimidazole (DRB), and the level of the isoforms evaluated over time. V2 of ATF4 was relatively stable (half-life 4.7 h), with a slow decline in levels over 8 hours. In contrast, V1 was extremely unstable (half-life 20 min) (Fig. 3d). Almost all V1 mRNA was gone by 1-hour post-treatment; this rapid decline is similar to the unstable *CMYC* transcript observed in these cells after PRMT5i treatment (Suppl. Fig. 3c). The difference in the stability of V1 and V2 was not dependent on PRMT5 activity, as PRMT5 inhibition did not change the transcript decay rates, only the relative proportion of the two transcripts (Fig. 3d). We also confirmed differential ATF4 transcript stability in another non-AML cell line (Suppl. Fig. 3d). Collectively, although we cannot exclude possible posttranscriptional conversion of V1 to V2 upon DRB treatment, it is likely that the rapid drop in V1 levels upon DRB treatment is transcript stability-related, as DRB also inhibits kinase activity essential for the splicing process (33).

The inherently unstable behavior of ATF4 V1 could be due to nonsense-mediated decay (NMD). NMD-dependent degradation was reported to lead to a decrease in the levels of several other alternatively spliced transcripts upon PRMT5 inhibition (10). To address this possibility, we performed knockdown of NMD regulator Upstream Frameshift 1 (UPF1) and found that it resulted in increased levels of ATF4 V2 but not the V1 mRNA (Fig. 3e, Suppl. Fig. 4a), thus confirming the previously reported role of NMD in the degradation of ATF4 V2 (34, 35). (Fig. 3e). In contrast, degradation of the unstable V1 did not rely on NMD as UPF1 knockdown did not affect its levels (Fig. 3e), indicating that NMD is not likely responsible for the unstable nature of V1. We also confirmed that PRMT5 inhibition and knockdown both downregulated V2 levels and upregulated V1 (Fig. 3e, Suppl. Fig 4a). Conversely, overexpression of PRMT5 increased V2 transcript levels and slightly decreased V1 (Suppl. Fig. 4b) in UCSD-AML-1 cells.

The absence of NMD dependent regulation of V1 indicates that this transcript may be detained in the nucleus to undergo degradation that is translation-independent. Approximately a third of human and mouse transcripts contain at least one intron, and many of these intron containing mRNAs (detained intron transcripts) remain in the nucleus (36, 37). Thus, we performed nuclear-cytoplasmic fractionation and investigated the abundance of V1 and V2 ATF4 mRNAs. The V1 form was predominantly found in the nucleus and its levels were further elevated in the nucleus following inhibition of PRMT5 (Fig. 3f). On the other hand, while the V2 form was more abundant in general, it was primarily found in the cytoplasm, and in contrast to V1 mRNA, its levels were decreased following PRMT5 inhibition (Fig. 3g). Ribosomal RPLP0 and small nucleolar RNA ACA19 were used as the fractionation controls (Fig. 3h). Together, these results suggest that PRMT5 catalytic activity is required for ATF4 splicing, generating the stable cytoplasmic V2 transcript of ATF4, while suppressing the unstable nuclear-detained V1 splice form containing the retained intron.

### PRMT5 inhibition results in reduced levels of ATF4 protein independently of P53 status

Our finding of PRMT5 inhibition leading to the production of unstable ATF4 V1 mRNA and lower levels of cytoplasmic V2 ATF4 mRNA suggested that ATF4 protein levels would be decreased. In preparation for these experiments, we observed increases in ATF4 protein levels upon higher cell density which is consistent with ATF4’s role in cell metabolism under stress conditions (26, 27, 38). Therefore, to avoid cell density and metabolic status-dependent effects, all ATF4 protein detection experiments were performed under cell number/density matched conditions. In keeping with the hypothesis above, treatment of cells with both PRMT5 inhibitors, but not their negative control compounds, effectively reduced ATF4 protein levels (Fig. 4a,b Suppl. Fig. 4c). Symmetric dimethylation of a well-characterized PRMT5 substrate SmBB’ was reduced, indicating the effectiveness of the inhibitors as previously reported (20, 21). Furthermore, decreasing amounts of ATF4 mRNA and protein, were associated with proportionately reduced levels of the ATF4 target phosphoserine aminotransferase 1 (PSAT1) (Fig. 4b). P53 protein levels were increased upon PRMT5 inhibition as reported before (7) (Fig. 4a), however, ATF4 downregulation was not dependent on p53 status. Wild type and *TP53* knockout HCT116 colon cancer cells, as well as *TP53* null K562 leukemia cells or *TP53* mutant 80309 EVI1 leukemia cells, all showed a reduction of ATF4 protein levels upon PRMT5 inhibition (Suppl. Fig. 4d-e).

**Fig. 4.**
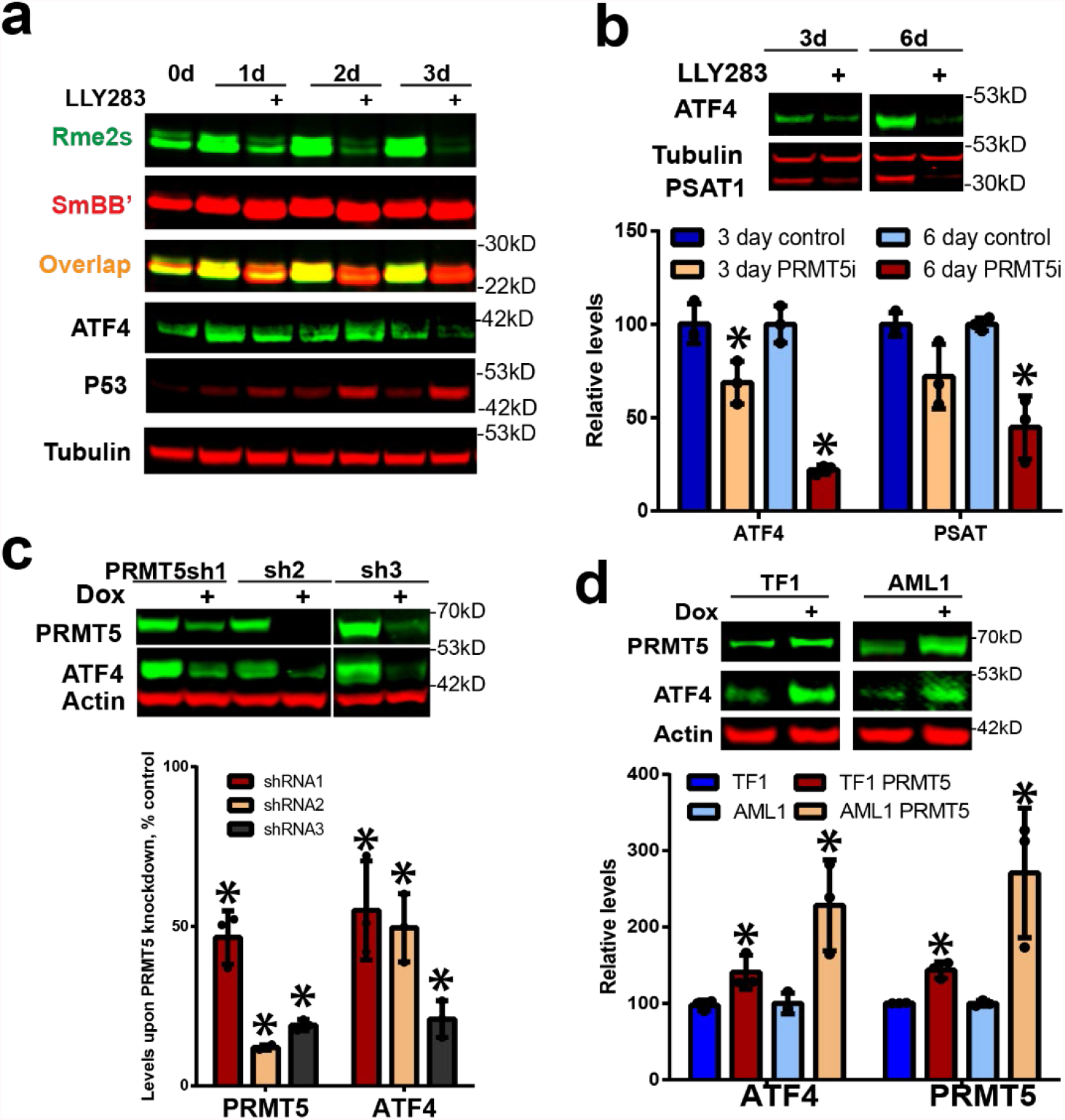
PRMT5 inhibition or knockdown reduces ATF4 protein levels. **a)** PRMT5 inhibitor LLY283 (100 nM) effectively reduces SmBB’ symmetric arginine dimethylation and ATF4 protein levels while it increases the p53 protein levels in OCI-AML-20 cells. **b)** ATF4 and its target PSAT1 protein levels were downregulated in response to 3 and 6 days LLY283 (100 nM) treatment, quantitation is presented in the graph below, N=3, mean±SD, * p<0.05 relative to the matched respective control. **c)** PRMT5 knockdown also downregulates the protein levels of ATF4, quantitation relative to control is presented in the graph below, N=3, * p<0.05, mean±SD. **d)** PRMT5 overexpression increased ATF4 protein levels. EVI1-high TF1 and UCSD-AML-1 leukemia cells were transduced with doxycycline (dox) inducible PRMT5, followed by PRMT5 and ATF4 protein levels assessment, two days after induction, quantitation is below; N=3, mean±SD, * p<0.05 relative to the matched respective control.

As PRMT5 can also regulate translation (39, 40), we investigated if PRMT5 inhibition would affect protein translation. For these experiments we used a reporter system (41) in which green fluorescent protein (GFP) is fused to the *ATF4* V2 upstream open reading frame (uORF), uORF1 and uORF2 elements controlling the translation of ATF4. uORF2 overlaps with the main productive ORF of ATF4 (see schematic in Suppl. Fig. 4f). Under baseline conditions, uORF2 is translated, thus preventing the translation of the main ORF. However, upon stress, the ribosomal scanning is impaired, bypassing the inhibitory uORF2 and resulting in translation and rapid increase in ATF4 protein levels from variant 2 main coding ORF (42). In this reporter system (41), ATF4 translation was not affected by PRMT5 inhibition (Suppl. Fig. 4f).

Additionally, we confirmed that similarly to PRMT5 inhibition, the genetic knockdown of PRMT5 by three different shRNAs also resulted in decreased levels of ATF4 (Fig. 4c). In contrast, overexpression of PRMT5 in two leukemia cell lines resulted in increased levels of ATF4 (Fig. 4d). Overall, these data indicate that PRMT5 catalytic activity regulates ATF4 splice variant switch, ATF4 protein levels, and downstream transcriptional program.

### PRMT5 regulates ATF4 driven redox balance in cells

ATF4 protein levels are increased in response to various stress stimuli (42, 43). Thus, we investigated if PRMT5 inhibition would result in a reduced capacity to induce ATF4 in response to stress. Treatment with several stress-inducing agents known to upregulate ATF4 protein levels (43, 44) consistently increased the levels of ATF4 protein (Fig. 5a) in OCI-AML-20 and UCSD-AML-1 cell lines. However, pretreatment of the cells with a PRMT5 inhibitor or PRMT5 knockdown attenuated the induction of ATF4 by the same stress inducers (Fig. 5a, Suppl. Fig. 5a). We also determined that while inhibition of PRMT5 affected ATF4, it did not regulated the pathway upstream of ATF4, phosphorylated eukaryotic initiation factor 2 (pEIF2) (Suppl. Fig. 5b) or the parallel UPR pathways of XBP1 and inositol-requiring enzyme 1 (IRE1) (Suppl. Fig. 5c-d).

**Fig. 5.**
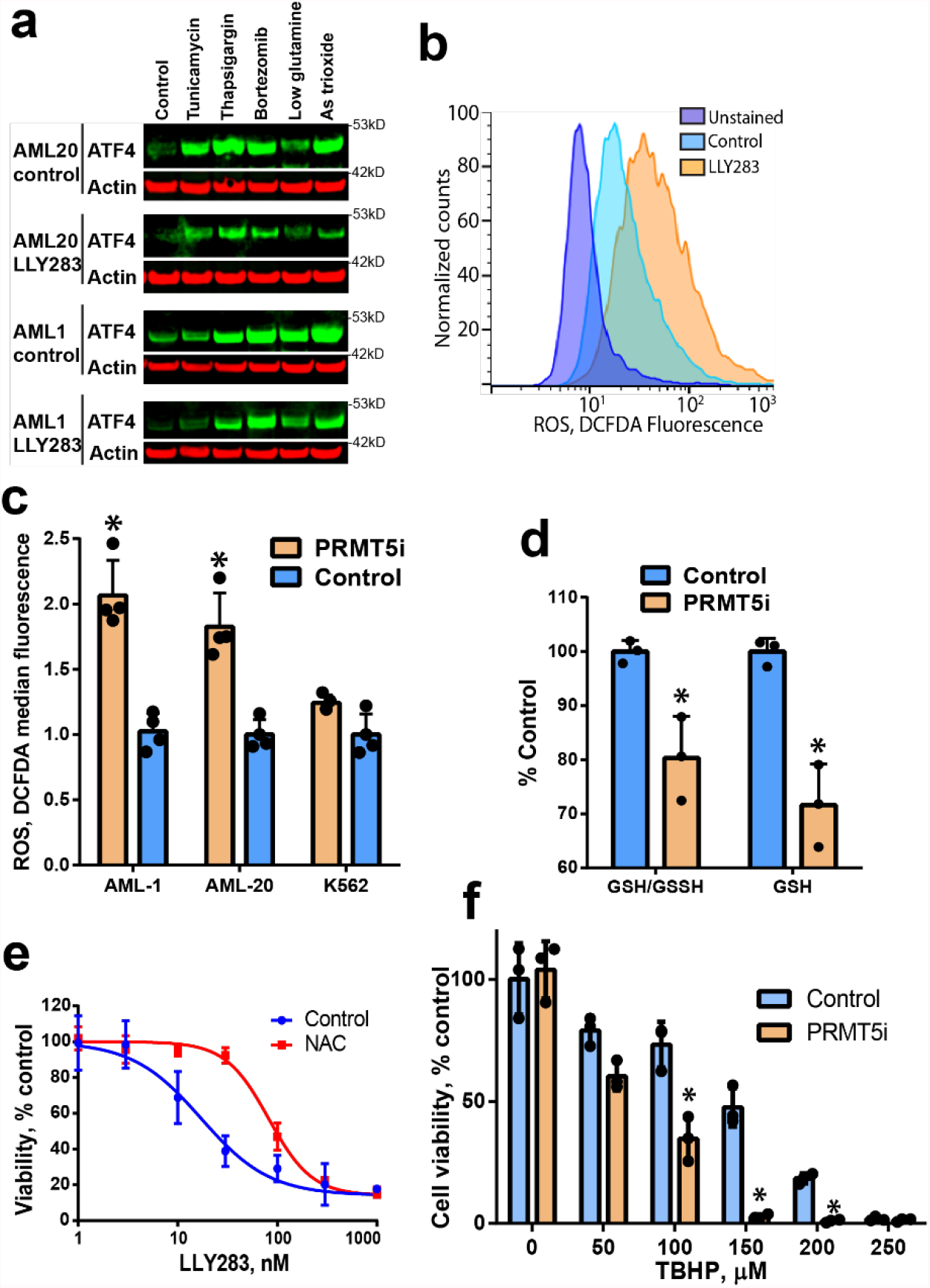
PRMT5 regulates ATF4 and downstream oxidative stress response. **a)** PRMT5 inhibition attenuates stressor-induced ATF4 levels. OCI-AML-20 and UCSD-AML-1 cells were pretreated with LLY-283 (100 nM) and exposed to ER stressors (8 h): the endoplasmic reticulum stressors tunicamycin (glycosylation inhibitor), thapsigargin (Ca^2+^ pump inhibitor), proteasome inhibitor bortezomib, glutamine starvation, or oxidative stress-inducing arsenic (As) trioxide. **b)** PRMT5 inhibition leads to higher ROS levels. Flow cytometry histograms for oxidative stress indicator DCFDA fluorescence in UCSD-AML-1 cells treated with 100 nM of LLY283 for four days. **c)** Quantitation of oxidative stress upon PRMT5 inhibition as in b) in UCSD-AML-1, OCI-AML-20, and K562 cells. N=4, means±SD, * p<0.05. **d)** GSH content is decreased upon PRMT5 inhibition (LLY283, 100 nM, 4 days) in UCSD-AML-1 cells. Absolute and GSSH normalized GSH levels are shown. N=3, means±SD * p<0.05. **e)** N-acetyl cysteine (NAC) antioxidant partially rescues PRMT5 inhibition-induced cell proliferation block in UCSD-AML-1. Cells were treated with LLY283 and (0.5 mM) NAC for four days. Cell viability was determined using resazurin assay N=3, means±SEM shown. **f)** PRMT5 inhibition (LLY238 30 nM 3 days) sensitizes UCSD-AML-1 cells to tert-butyl hydroperoxide (TBHP) exposure for 24 h. Cell viability was determined using resazurin assay, N=3, means±SD, * p<0.05.

ATF4 is induced in response to oxidative stress, and plays a critical role in cellular redox balance (45). As the above data showed attenuation of ATF4 upon stress and gene expression data indicated that antioxidant glutathione (GSH) pathway genes were downregulated by PRMT5 inhibition (Fig. 2), we investigated the levels of reactive oxygen species (ROS) using flow cytometry-based detection of oxidized 2’-7’-dichlorodihydrofluorescein diacetate (DCFDA). Treatment of EVI1-high OCI-AML-20 and UCSD-AML-1 cells with a PRMT5 inhibitor resulted in increased DCFDA fluorescence, indicating elevated ROS levels upon PRMT5 inhibition (Fig. 5b-c). Interestingly K562 cells that already have a high baseline ATF4 protein and do not overexpress EVI1 did not show increased levels of ROS (Fig. 5c, Suppl. Fig. 4e). In keeping with the higher levels of ROS in PRMT5 inhibited UCSD-AML-1, there was a decrease in levels of reduced GSH (Fig. 5d). Treatment of UCSD-AML-1 with the antioxidant agent N-acetyl cysteine (NAC) partially rescued PRMT5 elicited cell growth defect (Fig. 5e), indicating that oxidative stress contributes to the impairment of proliferation observed following PRMT5 inhibition. To investigate if PRMT5 inhibition would sensitize cells to oxidative stress, we treated cells with the PRMT5 inhibitor LLY283 for 3 days and exposed them to tert-butyl hydroperoxide (TBHP) for 24 h. When PRMT5 was inhibited, UCSD-AML-1 cells displayed reduced cell viability in response to TBHP as well as known oxidative stress inducers such as arsenic trioxide and cisplatin (Fig. 5f, Suppl Fig. 5e,f). In addition, overexpression of doxycycline-inducible ATF4 attenuated LLY283-mediated cell proliferation inhibition (Suppl. Fig. 5g). Taken together, PRMT5i sensitized cells to oxidative stress, while NAC antioxidant or ATF4 overexpression attenuated the PRMT5i-induced cell proliferation defect, demonstrating that the PRMT5-ATF4 axis plays a role in regulating cellular redox state.

### EVI1 sensitizes cells to PRMT5 inhibition by reinforcing low baseline levels of ATF4

Finally, to understand the mechanism behind the PRMT5 dependence of EVI1 overexpressing cells (Fig.1a), we investigated whether EVI1 overexpression would sensitize cells to PRMT5i. Overexpression of EVI1 in OCI-AML5 leukemia cells that are *TP53* wild type and have very low endogenous EVI1 resulted in greater impairment of cell proliferation following PRMT5 inhibition (Fig 6a). To determine the cellular pathways affected by EVI1 that may explain sensitivity to PRMT5 inhibition, we used Gene Set Enrichment Analysis (GSEA). GSEA of the mouse bone marrow lineage negative cells infected with a retroviral construct of EVI1 (GSE34729 dataset) (46) showed a significant decrease in the ATF4 regulated gene signature relative to the control cells (Fig. 6b). As the mRNA levels of ATF4 were not altered by EVI1 overexpression in the GSE34729 data set and we found no change in ATF4 mRNA levels upon overexpression of EVI1 (Suppl. Fig. 6a), we wondered if EVI1 regulated the protein levels of ATF4. We opted to overexpress EVI1 in very low endogenous EVI1 OCI-AML5 leukemia cells (used above) and additional pro-monocytic U937 cell line (*TP53* mutant) that has been used in previous EVI1 studies (47). In both cell lines overexpression of EVI1 resulted in downregulation of protein levels of ATF4 (Fig. 6c). To support the ectopic EVI1 overexpression studies, we also utilized EVI1 knockdowns in relevant inv(3) EVI1-high USCD-AML-1 cells. The knockdown of EVI1 resulted in elevated ATF4 protein levels (Fig. 6d, Suppl. Fig 6b). To further understand the mechanism behind EVI1 downregulation of ATF4 protein, we asked if ATF4 translation or protein stability were affected by EVI1. The ATF4 reporter system indicated no translational regulation of ATF4 by EVI1 (Suppl. Fig. 6c), and this was consistent with the lack of EVI1 driven regulation of the PI3K/Akt pathway in OCI-AML5 cells (Suppl. Fig. 6d). On the other hand, EVI1 overexpression led to higher levels of ATF4 ubiquitylation and more rapid ATF4 protein degradation (Suppl. Fig. 7a, b). Notably, EVI1 overexpression or PRMT5i did not affect the levels of another critical oxidative stress response regulator, NRF2 (Suppl. Fig. 7c), indicating that responses seen were due to ATF4.

**Fig. 6.**
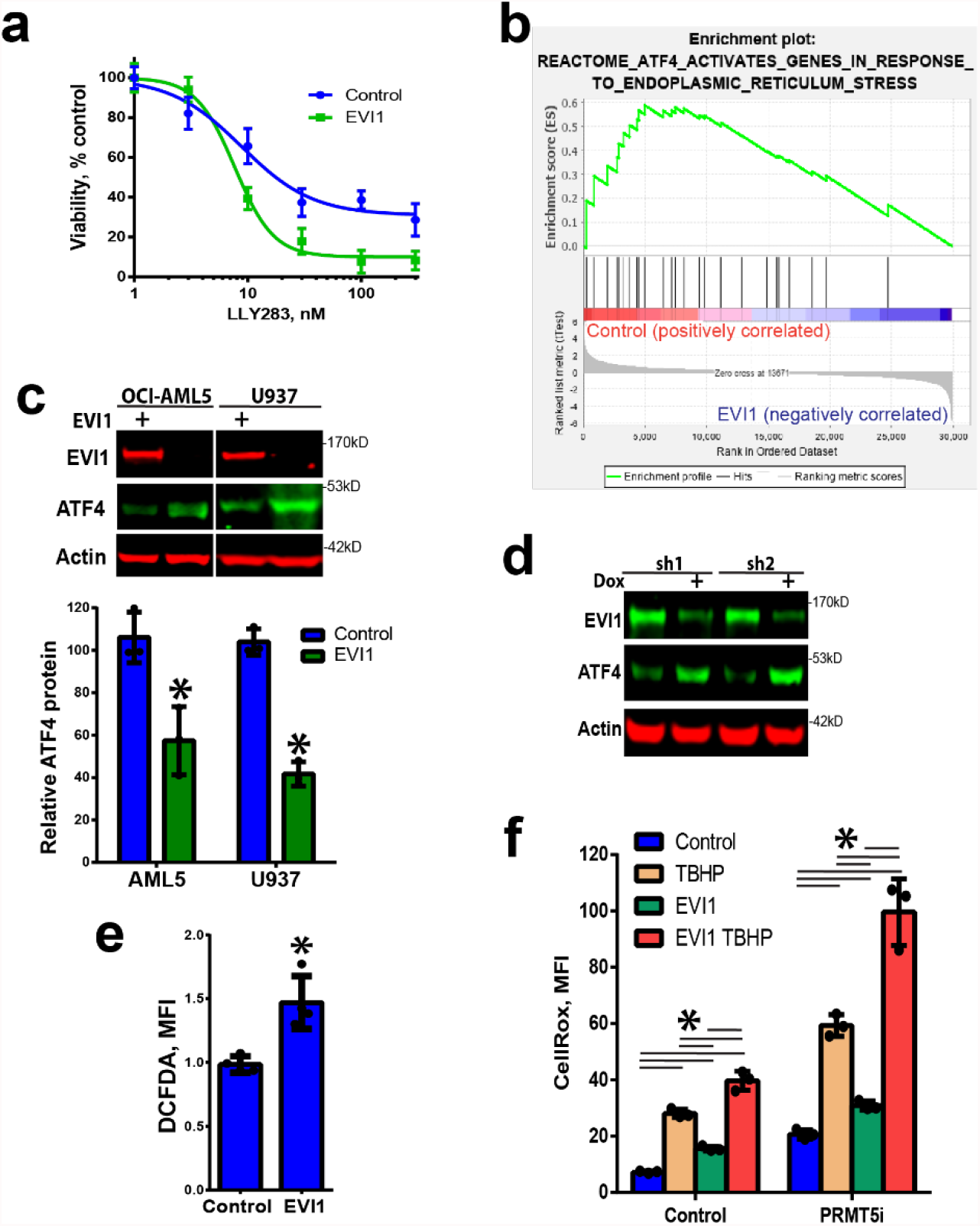
EVI1 overexpression results in decreased ATF4 levels and sensitization to PRMT5 inhibition. **a)** EVI1 overexpression in OCI-AML5 cells sensitizes to PRMT5 inhibition. Cell viability was assessed by resazurin assay after 5 days (N=3, means±SEM). **b)** Gene set enrichment analysis (GSEA) of EVI1 overexpressing cells (GSE34729) indicates significantly decreased enrichment for the ATF4 signature (negative correlation between EVI1 and ATF4 signatures NES=1.67, p=0.01). **c)** EVI1 overexpression leads to lower ATF4 protein levels in U937 and OCI-AML5 cells (no endogenous EVI1 expression), quantitation below (N=3, * p<0.05, means±SD). **d)** EVI1 knockdown results in increased ATF4 levels in UCSD-AML-1 cells (EVI1-high). Dox indicates doxycycline induction of shRNA (2 days). **e)** Increased oxidative stress in U937 cells overexpressing EVI1 (N=4, means±SD, * p<0.05). **f)** Increased oxidative stress in AML5 cells overexpressing EVI1 is potentiated by PRMT5 inhibition (LLY283, 100 nM, 4 days) and TBHP (200μM, 2h), N=3, means±SD, * p<0.05 two-way ANOVA, Holm-Sidak multiple comparison test.

**Fig. 7.**
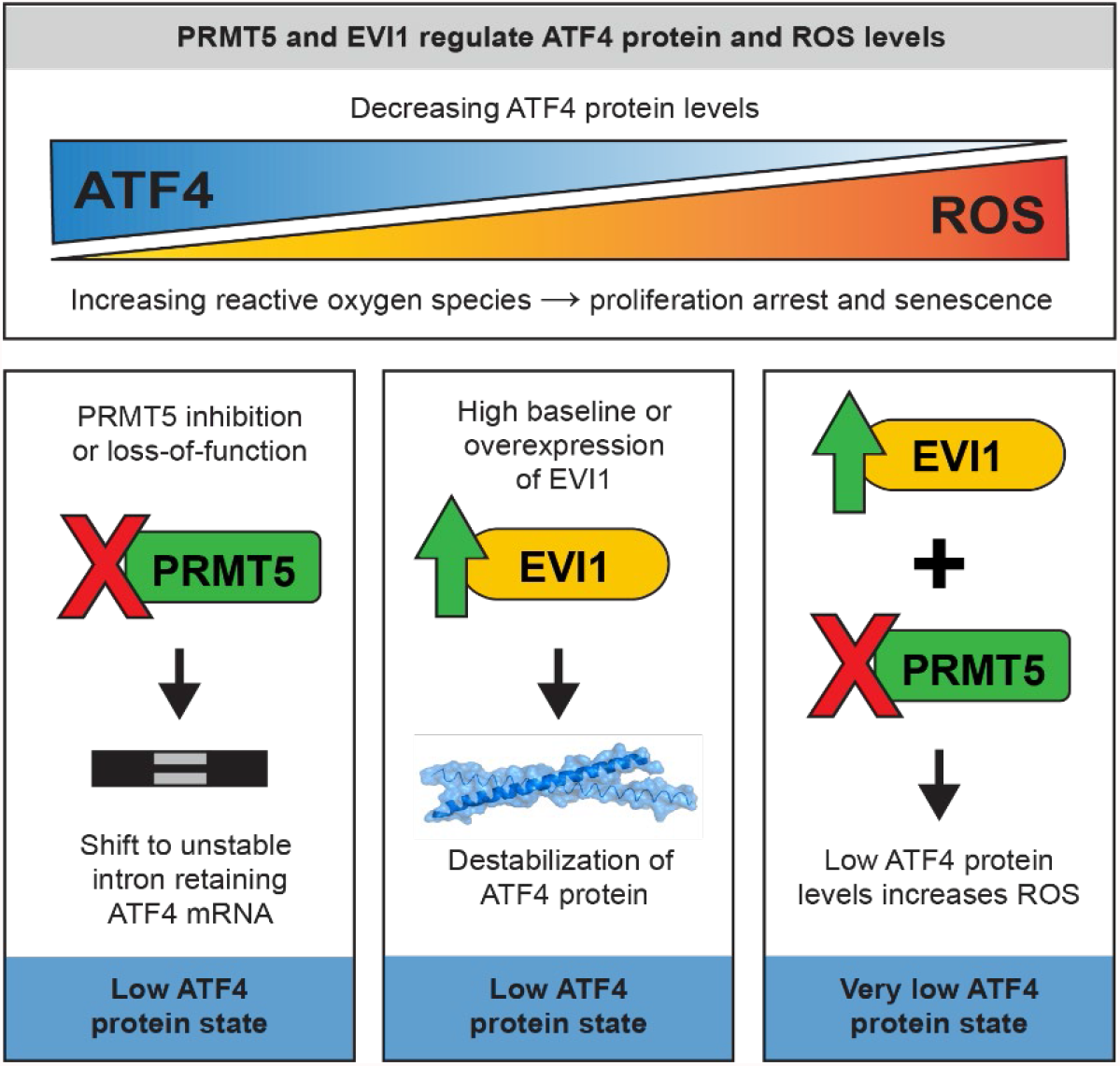
A schematic of ROS and ATF4 regulation by PRMT5 and EVI1. PRMT5 inhibition leads to unstable intron retaining ATF4 mRNA and low ATF4 protein levels, while overexpression of EVI1 also decreases ATF4 protein levels. Both, high EVI1 and PRMT5i result in a very low ATF4 state and increased intracellular ROS affecting cell proliferation, survival, and senescence.

To address the functional significance of EVI1 and PRMT5-ATF4 axis in oxidative stress, we investigated cellular ROS levels. The overexpression of EVI1 in U937 cells induced increased cellular ROS levels (Fig. 6e) that was consistent with the downregulation of ATF4 protein. Additional experiments using CellROX ROS indicator confirmed that EVI1 overexpression leads to oxidative stress in another leukemia cell line OCI-AML5 (Fig. 6f). Cells with overexpression of EVI1 and inhibition of PRMT5 displayed elevated levels of ROS that were further increased when PRMT5 was inhibited in EVI1 overexpressing cells and potentiation of ROS levels was observed with exposure to TBHP (Fig. 6f). Thus, oxidative stress in EVI1-high cells is further exacerbated by PRMT5 inhibition.

These results link EVI1 overexpression to the suppression of the ATF4 signature and increased oxidative stress. Taken together, our study indicates that the loss of PRMT5 catalytic activity results in altered splicing and downregulation of the ATF4 pathway leading to elevated cellular oxidative stress levels. Furthermore, this increased oxidative stress is exacerbated by EVI1 overexpression that also converges on ROS regulation, potentially creating a greater vulnerability to PRMT5i in EVI1-high cells (Fig 7).

## Discussion

Here we report a mechanism by which PRMT5 regulates ATF4 transcription factor and the oxidative stress response. The loss of PRMT5 catalytic activity led to intron retention and expression of the V1 form of ATF4 mRNA that is unstable and retained in the nucleus of dividing leukemia cells. The resulting decrease of cytoplasmic V2 mRNA form and ATF4 protein compromised the cellular oxidative stress response and resulted in elevated ROS. We also found that leukemia cells expressing the EVI1 oncogene showed a pronounced dependence on PRMT5. In cells expressing EVI1, ATF4 protein levels and downstream gene signature were suppressed, while ROS levels were elevated. Thus, PRMT5 ensures the fidelity of ATF4 transcript splicing, safeguarding the cellular redox balance critical in high oxidative stress states such as EVI1 overexpression.

PRMT5 depletion or inhibition leads to aberrant splicing and transcriptome changes (7, 8, 10, 30, 31). We found that PRMT5 inhibition or knockdown resulted in downregulation of ATF4 and downstream ATF4 regulated UPR genes. Interestingly, the ATF4 gene has an evolutionary conserved weak/non-consensus intron 1 splice donor site. Such weak splice donor sites are dependent on PRMT5 catalytic function (7). Compromised splicing at this site results in the intron retaining V1 isoform that we found to be unstable in dividing leukemia cells. It is possible that the stability of V1 is cell-type-specific as a study on ATF4 variant regulation reported high levels of the V1 mRNA in normal mature human leukocytes where V1 seems to be stable (48). Additionally, in granulocytic differentiation, widespread splicing factor alterations contribute to a distinct differentiation phenotype that is associated with intron retention in a set of genes, including ATF4 (49). Thus, splice forms of ATF4 may have functional importance in differentiation. A study in glioblastoma cells noted that PRMT5 inhibition results in intron retention and transcript nuclear detention, also linking it to differentiation (30). Taken together, we found that in dividing leukemia, V1 is extremely unstable and detained in the nucleus, thus explaining how the splicing isoform switch results in overall lower ATF4 transcript and protein levels.

PRMT5 function is essential for normal hematopoiesis (17), maintenance of chronic myelogenous leukemia stem cells (50), and leukemia with MLL rearrangements and splicing factor mutations (8, 10, 19). Our findings on PRMT5 inhibition leading to cell cycle arrest and senescence are consistent with previous reports in glioblastoma, AML, and other cancers (8, 30, 51-53). ATF4 is also required for normal hematopoietic system function and controls the UPR in HSCs and LSCs (54, 55). Previous studies indicated that ATF4 is a central regulator of cellular redox balance and oxidative stress (45), and ATF4-mediated attenuation of oxidative stress plays a role in erythropoiesis maintenance and HSC development in mice (55). We found that PRMT5 inhibition results in reduced levels of ATF4 mRNA and protein and increased levels of ROS. Interestingly, PRMT5 regulation of the DNA damage response has been noted in numerous studies focusing on splicing (14) or direct arginine methylation of DNA repair factors (56). Oxidative stress is a known contributor to DNA damage, and PRMT5 loss has been reported to induce oxidative DNA damage (18). However, the mechanism of the oxidative stress increase upon PRMT5 loss or inhibition was not clear. Our work shows that PRMT5 catalytic activity is required for maintaining ATF4 mRNA and protein levels and attenuation of oxidative stress, thus, providing important insight into PRMT5 function in cellular ROS regulation.

Interestingly, a PRMT5-mediated transcriptional mechanism has been linked to antioxidant gene regulation and chemotherapy-induced formation of a β-catenin-JDP2-PRMT5 complex (57). Upon genotoxic stress, PRMT5 associates with β-catenin and JDP-2 to drive transcription of GSH metabolic genes. In keeping with the antioxidative function of PRMT5, knockdown of PRMT5 in ovarian cancer cells increased levels of ROS (57). Oxidative stress has also been reported to lead to PRMT5-mediated increase in H4R3 symmetric dimethylation and the base excision repair complex assembly that ensured oxidative DNA lesion repair (58). However, a study in renal cells subjected to ischemia-reperfusion stress found that inhibition of PRMT5 ameliorated pyrroptosis and oxidative stress (59) and glutathionylation was found to decrease PRMT5 activity (60). Thus, collectively oxidative stress control by the PRMT5-ATF4 axis is a novel mechanism that is a part of a more extensive network regulating cellular ROS physiology.

Oxidative stress is also likely to contribute to the senescence phenotype we observe with prolonged inhibition of PRMT5 as oxidative injury is a well-known causative factor for cellular senescence (61, 62). PRMT5 loss of function has been reported to elicit the senescence response in glioblastoma and osteosarcoma cells (51, 63). In contrast, ATF4 was found to antagonize cellular senescence (64). Although in general, senescence is considered beneficial for the control of cancer cell proliferation, several studies indicated that chemotherapy-induced senescence can lead to proliferation arrest escape and the emergence of aggressive drug resistant cell population in AML and other malignancies (65, 66). Thus, while PRMT5 inhibitor-based therapeutics are promising candidates in several cancers, the complex outcomes of PRMT5 inhibition and implications of cell senescence on long term response need to be considered, especially in rational selection of drug combinations.

Leukemia and MDS with splicing factor mutations or MLL translocations display dependency on PRMT5 function (8, 10, 15, 19, 67). We found that leukemia cells with EVI1 overexpression also had increased sensitivity to PRMT5 inhibition and reduced levels of ATF4 protein. The EVI1 proto-oncogene is a transcription factor (25) that regulates various transcriptional programs (46, 68-74), including an extensive reprogramming of metabolic pathways (75). Although EVI1 has been shown to downregulate PTEN (76), we did not find that EVI1 overexpression affected PI3K/AKT pathway. The latter is known to control the levels of cellular ATF4 (77-79). Instead, ATF4 protein stability was decreased by EVI1 overexpression. As the degradation of ATF4 has been associated with several kinases and signaling pathways (80-82), these pathways may be influenced by EVI1.

The reported oxidative stress upregulation by EVI1 (83, 84) is consistent with our findings of the antagonistic relationship between EVI1 and ATF4. Through increased oxidative stress, EVI1 creates a dependency on PRMT5 function in maintaining a cellular redox state that is critical for cell survival. In a broader sense, excessive production of ROS is a common feature of leukemic and other cancer cells driven by several oncogenes, including EVI1 (85). Consequently, future studies are needed to determine links between high basal levels of oxidative stress and dependence on PRMT5 function in other cancers as well as their therapeutic relevance.

## Conclusions

Our work sheds light on the role of PRMT5 regulated splicing in oxidative stress. PRMT5 inhibition results in the splicing switch from the stable cytoplasmic ATF4 mRNA variant to the unstable intron-retaining, nuclear-localized variant. The subsequent reduction in ATF4 transcript and protein levels leads to downregulation of the ATF4 transcriptional program, elevated oxidative stress, and cell senescence. ATF4 protein levels are also suppressed by the EVI1 oncogenic transcription factor contributing to a high ROS environment in EVI1 overexpressing cells. Subsequently, these cells display higher sensitivity to PRMT5 loss of function. Hence, we identified a functional link between PRMT5 and EVI1, where both converge on the ATF4 oxidative stress pathway.

## Materials and methods

### Cell culture and treatments

Patient sample protocols for this study were approved by the University Health Network’s Research Ethics Board. Patient samples were collected with informed consent in accordance with the procedures outlined by the University Health Network’s Research Ethics Board approval. All biological samples were collected with informed consent according to procedures approved by the Research Ethics Board of the University Health Network (UHN; REB# 01-0573-C) and viably frozen in the Leukemia Tissue Bank at Princess Margaret Cancer Centre/ University Health Network. Cryopreserved, ficoll purified patient mononuclear cells from either bone marrow or peripheral blood were rapidly thawed, washed and cultured in IMDM media (Gibco/Wisent) with 2 mM L-glutamine, 10% fetal bovine serum (FBS) (Wisent), 55 uM β-mercaptoethanol (Gibco), 100 ug/mL Primocin (InvivoGen), 100 ng/mL SCF, 50 ng/mL FLT3L, 40 ng/mL THPO, 20 ng/mL IL3, and 20 ng/mL GM-CSF (Shenandoah Biotech or Custom Biologics) in 6-well plates (Greinier) seeded with confluent OP9. OP9 mouse stromal cells (ATTC) were previously cultured in α-MEM media with GlutaMAX (Gibco) containing 20% FBS (Wisent), 55 µM β-mercaptoethanol (Gibco) and 100U/100 ug/mL penicillin/streptomycin (Gibco) at 37 °C/5% CO2. Patient cells were allowed to recover from thawing for 2-3 days prior to subsequent experiments and topped with fresh media or transferred to new OP9-containing plates as necessary.

Cell lines were cultured according to standard aseptic mammalian tissue culture protocols in 5% CO2 at 37 °C with regular testing for mycoplasma contamination using the MycoAlertTM Mycoplasma Detection Kit (Lonza). HCT116, HEK293T, cells were cultured in DMEM supplemented with 10% FBS (Wisent) and 100U/100 μg/mL pencillin/streptomycin (Wisent). UCSD-AML-1, OCI-AML5, U9376 were cultured in IMDM media with GlutaMAX (Invitrogen) containing 10% FBS (Wisent), and 10 ng/ml GM-CSF (Shenandoah Biotech or Custom Biologics) and 100U/100 μg/mL penicillin/streptomycin (Wisent). K562 cells were grown in the above media without GM-CSF.

PRMT5 inhibitors GSK591 and LLY283 as well as their negative controls SGC2096, LLY284 were from the SGC probe collection(86).

5,6-dichloro-1-beta-D-ribofuranosylbenzimidazole (DRB) was from Sigma and was used at 100 µM for 2h. Cells were exposed for 1-18h to tunicamycin (Sigma, 1 mg/ml), thapsigargin (Sigma, 300 nM), bortezomib (Cayman 100 nM), arsenic trioxide (Sigma, 10 μM) (see figure legends for exposure time).

### Antibodies

Anti-Rme2s (#13222), ATF4 (#1185), XBP1 (#12782), IREa (3294), EIF2A (#9722), pEIF2A (#3398), UPF1 (#12040), phospho p90RSK, pAKT. P44/42 MAPK, pS6 (#7100), NRF2 (#12721), HA (#3724) were purchased from Cell Signaling Technologies. Antibodies for PRMT5 (#ab109451), p53 (#ab1101), H3 (#10799) and β-actin (#ab3280) were purchased form Abcam. Anti-Flag (#F4799) was from Sigma. Anti-SmBB’ (#sc-130670) and Anti-B-tubulin (sc-5274) were purchased from Santa Cruz Biotechnologies. Anti-PSAT1(PSAT1-2-s) was purchased from Developmental Studies Hybridoma Bank. Secondary goat-anti rabbit-IR800 (#926-32211) and donkey anti-mouse-IR680 (#926-68072) were purchased from LiCor.

### Constructs, transduction and knockdown

Flag tag EVI1 lentiviral construct was a gift from Dr L. Salmena (University of Toronto) (87). Lentiviral constructs were packaged and transduced as described before (88). Briefly, third generation packaging plasmids pRSV-Rev (Addgene 12253), pMDLg/pRRE (Addgene 12251), pCMV VSV-G (Addgene 8454), and the lentiviral vector were transfected into HEK293T and viral supernatants harvested at day 3. ATF4 reporter was a gift from Dr J.E. Dick (University of Toronto), (Addgene #155032) (54). ATF4 cDNA was cloned into a doxycycline inducible pLVX (Clontech) vector, virus produced and transduced as above. PRMT5, ATF4, and EVI1 were cloned into a derivative of pSTV doxycycline inducible lentiviral vector (Dr A.C. Gingras). Additionally, EVI1 was cloned into pSMALGFP vector (Addgene #161785). Ubiquitin with HA tag vector was from Addgene (#18712).

PRMT5 shRNAs in pLKOpuro vector were from Sigma sh1 (H3) sequence (CCTCAAGAACTCCCTGGAATA), sh2 (H4) (GCCCAGTTTGAGATGCCTTAT), and sh3 (H5) (GGCTCAAGCCACCAATCTATG). EVI1 shRNAs were cloned into pLKO2Tetpuro (Addgene #21915) sh1 (TGCAGGGTCACTCATCTAAAG), sh2 (GCACTACGTCTTCCTTAAATA), UPF1 sh (AGATATGCCTGCGGTACAAAG).

### Western blotting and immunoprecipitation

Cells were lysed in lysis buffer (20 mM Tris-HCl pH 8, 150 mM NaCl, 1 mM EDTA, 10 mM MgCl2, 0.5% TritonX-100, 12.5 U/ml benzonase (Sigma), complete EDTA-free protease inhibitor cocktail (Roche)). After 30 sec incubation at RT, SDS was added to final 1% concentration and proteins were quantitated by BCA Protein Assay Kit (Pierce). Total cell lysates were resolved in 4-12% Bis-Tris Protein Gels (Invitrogen) and transferred in for 1.5 h (80 V) onto PVDF membrane (Millipore) in Tris-Glycine transfer buffer containing 20% methanol and 0.05% SDS. Blots were blocked for 1 h in blocking buffer (5% milk in PBS) and incubated with primary antibodies in blocking buffer (5% BSA in PBST: 0.1% Tween 20 PBS) overnight at 4 °C. After five washes with PBST the blots were incubated with goat-anti rabbit (IR800) and donkey anti-mouse (IR 680) antibodies in Odyssey Blocking Buffer (LiCor) for 1 h at RT and washed five times with PBST. The signal was read on an Odyssey scanner (LiCor) at 800 nm and 700 nm. Band intensities for western blot analysis were determined using Image Studio Ver 5.2 (Licor). Unless otherwise noted, all immunoblotting images were representative of at least three biological replicates from independent experiments. For immunoprecipitation experiments HEK293 cells were transfected with HA ubiquitin, ATF4 (Flag tagged) and EVI1 or empty pSTV. Cells were lysed in the above lysis buffer, lysates spun down and precipitated overnight with ATF4 or Flag antibodies. The precipitated complexes were captured by protein G magnetic beads (BioRad) and washed 3 times with lysis buffer with subsequent SDS PAGE separation and blotting with an anti-HA antibody.

### Cell viability assessment

Patient cells were transferred to GFP labelled OP9 cells in 96-well plates for compound testing at a density of either 25000 or 50000 cells per well in 100 µL media with 0.1% DMSO/compound. OP9 cells were transduced with GFP lentivirus (derivative of pRRL-GFP vector from Dr Trono group, Addgene 12252). OP9 labelling with GFP allowed for differentiation of OP9 cells from leukemia cells in flow cytometry experiment. On day 3-4, an additional 100 uL of media and compound were added. Viable cell number was assessed at day 6 for PRMT5 inhibitor GSK591 ranging from 0 to 2 µM, or negative control (SGC2096) treatments. To assess viability, cells were transferred to 96-well round well suspension plate (Starstedt) along with trypsinized (30 uL 0.25% Trypsin-EDTA; Wisent) OP9 stroma and attached leukemic cells. Cells were resuspended in PBS (Wisent) with 2% FBS and 0.2 µM Sytox Blue (Life Technologies) viability dye. A total of 50 μL suspension of cells for each well were run through a MACSQuant VYB (Miltenyi) flow cytometer and MACSQuantify software was used to determine viable leukemic cell number (GFP negative, Sytox blue negative cells).

OCI-AML-20 line, 80309, 120781, or UCSD-AML-1 line were transduced with GFP or mCherry lentivirus and cells FACS sorted to obtain a homogenous GFP/mCherry-expressing population. OCI-AML-20 line, 80309, 120781 cells were maintained on confluent monolayer of OP9 bone marrow stromal cells. Cells were seeded at density of 2000-5000 cells per well of a 96 well plate. Proliferation was monitored by counting GFP/mCherry-expressing cells over time using an IncuCyte live-cell imaging and analysis platform (Sartorius).

For the apoptosis assay, non-GFP cells as above were seeded and treated with PRMT5 inhibitors and Caspase-3/7 Green Dye for Apoptosis (Essen Bioscience) that couples the activated caspase-3/7 recognition motif (DEVD) to a DNA intercalating dye to enable quantification of apoptosis over 6 day timeline.

UCSD-AML-1 and OCI-AML5 cell viability was determined using resazurin assay (Sigma) as per manufacturer’s instructions. Briefly, resazurin at final concentration 0.01 mg/ml was added to cells and fluorescent resorufin signal was measured using a plate reader at excitation/emission 570/585 nm.

### S-phase, quiescence, senescence, and reporter assays

S-phase was measured using an EdU assay. Control, 1µM PRMT5 inhibitor or negative control treated (3 or 6 day) OCI-AML-20 cells were incubated with 10 µM EdU (Invitrogen) for 2 h at 37 °C. Labelled cells were fixed with 4% PFA and Click-IT Plus Alexa Fluor 488 flow cytometry kit (Invitrogen) was used according to the manufacturer’s instructions. The cells were processed using a MACSQuant VYB flow cytometer (Miltenyi) and flow data was analysed with FlowLogic software. Initial gating by FSC-A and SSC-A was used to remove debris. Singlet gates were determined by FSC-A/FSC-H. The cell number was recorded. Alexa Fluor 488 signal was used to identify S phase cells.

Quiescent cells were measured as previously described(89). In brief, ∼250-500k control or treated OCI-AML-20 cells were incubated with 10 µg/mL Hoechst 33342 at 37°C for one hour. A final concentration of 0.5 µg/mL pyronin Y (Sigma) was added for additional 15-20 minutes at 37°C. Cells were put on ice and analysed using the same instrumentation and software as EdU assay. Gating strategy was similar as above except pyronin Y and Hoechst signals were used to identify G1, G0, G2/M and S phase populations.

Senescence assay utilizing β-galactosidase was performed as described(90). Briefly OCI-AML-20 cells were spun down and washed with PBS, fixed with 2% formaldehyde and 0.2% glutaraldehyde for 5 minutes, washed twice with PBS and incubated in the staining solution containing 40 mM citric acid/Na phosphate buffer, 5 mM K4[Fe(CN)6] 3H2O, 5 mM K3[Fe(CN)6], 150 mM sodium chloride, 2 mM magnesium chloride and 1 mg/ml X-gal for 18 hours. Cells were imaged with EVOS microscope (Invitrogen). Cells positive for β-galactosidase were scored from at least 50 cells in 3-4 fields in biological replicates.

ATF4 reporter assays utilized pSMALB construct (54). (gift for Dr J. Dick, Addgene #155032) that was packaged and transduced as above. The GFP/BFP intensities were measured by flow cytometry and the ratio was assessed described.

### RNAseq analysis

OCI-AML-20 cells were treated with 1 µM GSK591 for six days; previous studies showed complete PRMT5 inhibition by GSK591. RNA was extracted using TRIzol (Invitrogen) and quality was assessed using a Bioanalyzer (Agilent Technologies). Sample libraries were prepared using the Illumina TruSeq Stranded mRNA sample preparation kit. Sequencing was performed using Illumina Next-Seq500 using 75-cycle paired-end protocol and multiplexing at the Princess Margaret Genomics Centre.

RNA-Seq reads were aligned to the hg19 human reference genome using Bowtie(91) (2.0.5) and Tophat(92) (2.0.8) using default settings. Cufflinks(93) (2.1.1) was used to compute and normalise read counts and call differentially expressed genes. All the data for RNA-seq are available through GEO (GSE163305, raw data submission to EGA in progress).

Differential expressed genes were plotted using R Enhancedvolcano package (Kevin Blighe, Sharmila Rana and Myles Lewis, EnhancedVolcano: Publication-ready volcano plots with enhanced colouring and labeling. R package version 1.4.0. Available at: https://github.com/kevinblighe/EnhancedVolcano) in R version 3.6.1 (R Core Team (2019). R: A language and environment for statistical computing. R Foundation for Statistical Computing, Vienna, Austria. URL https://www.R-project.org/).

### Splicing and enrichment analysis

RNAseq alternative splicing events were quantified using the python package rmats-turbo version 4.1.0 ((94) and rmats-turbo: Xie Z., Xing Y., rMATS-turbo. http://rnaseq-mats.sourceforge.net/. Accessed 20 August 2020) with settings “t paired”, “readLength 75”, and “variable-read-length” enabled. Sashimi plots were generated with the rmats2sashimi python package(95) (version 2.0.2) and splicing event plots were generated with R maser package (version 1.4.0, Diogo F.T. Veiga (2019); maser: Mapping Alternative Splicing Events to pRoteins. R package version 1.4.0. https://github.com/DiogoVeiga/maser).

The GSEA tool was used to assess enrichment for gene signature(96). Gene set enrichment analysis for splicing events filtered through maser for FDR <0.05 were done using clusterProfiler(97) (version 3.14.3) in R package using supported pathways from the ReactomePA package(98) (version 1.30.0). Reactome pathway enrichment and transcription factor sites (TTRUST) categories were identified using Enrichr (99).

### Gene expression validation, ATF4 variant detection, and cell fractionation

RNA was extracted by TRIzol (Invitrogen) using the manufacturer’s protocol. Isolated RNA was reverse transcribed using the iScript gDNA clear reagents (Biorad) and an aliquot of the DNAse treated reaction was saved for the later PCR confirmation of the absence of genomic DNA. RT-qPCR was performed using a CFX384 Real-Time PCR Detection System (Bio-Rad) and SSoFAST SYBR real time PCR reagent (Biorad). Relative transcript levels were determined by the delta-delta Ct method and normalized to the housekeeping genes of 18s, TBP, and RPLP0. Primer sequences are shown in Supplementary Table 1.

For cell fractionation, cytoplasmic and nuclear fractions were isolated and RNA extracted using PARIS kit (ThermoFisher). The resulting RNA was DNase treated and cDNA generated as above. A fraction of DNase treated reaction was reserved for PCR to ensure that no genomic DNA contamination was present. PCR was performed with primers in Suppl. table 1. Absolute transcript quantitation was performed using DNA standards and quantitated in Maestro CFX software (BioRad).

### Oxidative stress, ROS and GSH/GSSG measurements

ROS was measured using 6-carboxy-2’,7’-dichlorodihydrofluorescein diacetate (H_2_DCFDA) or CellROX orange ThermoFisher Scientific) according to the manufacturer’s instructions. Cells were loaded with 10 μM H_2_DCFDA in PBS for 30-60min, recovered in cell media for 30 min and loaded and unloaded control cells analysed for FITC signal using flow cytometry as described above. Cells were loaded with 5 μM CellROX in media for 30 min and loaded and unloaded control cells analysed for mCherry channel signal using flow cytometry as described above GSH and GSSG levels were measured using GSH/GSSG-Glo assay kit from Promega according to the manufacturer’s instructions. Briefly in a 96 well (Corning) plate 20,000 cells were lysed with 25 μl GSH or GSSG lysis reagents and luciferin generation reagent was added (50 μl) followed by luciferin detection regent (100 μl). GSH and GSSG were used to generate standard curves and luminescence was measured using BMG Labtech ClarioStar plate reader.

### Statistical analyses

All measurements were taken from distinct samples/biological replicates. Statistical analyses were performed and plotted using Graphpad Prism 6. All data are presented as mean±standard deviation, unless otherwise stated. Values were obtained from at least three biological replicates and indicated in each respective figure legend or the figure. Statistical significance was determined by T test when comparing control and treated matched samples, analysis of variance (ANOVA) when normal distribution and equal variance was determined, followed by Tukey’s post-hoc test for multiple group comparisons. For non-normally distributed data, a non-parametric test (Kruskal-Wallis) was used, followed by multiple group comparisons using Dunns or Holm-Sidak’s tests. A p value of <0.05 was considered statistically significant. Statistical tests are indicated in the figure legends unless T test was used for control and treated sample comparison.

## Acknowledgements

We thank Dr L. Salmena and Dr J. Dick for providing reagents. We thank for the support of Leukemia Tissue Bank at Princess Margaret Cancer Centre / University Health Network as source for primary patient samples and clinical data. This work was supported by Leukemia Lymphoma Society of Canada, the Canadian Institutes of Health Research (FDN154328 to CHA), and a Natural Sciences and Engineering Research Council Postdoctoral Fellowship to DD and VV. The Structural Genomics Consortium is a registered charity (no: 1097737) that receives funds from AbbVie, Bayer AG, Boehringer Ingelheim, Genentech, Genome Canada through Ontario Genomics Institute [OGI-196], the EU and EFPIA through the Innovative Medicines Initiative 2 Joint Undertaking [EUbOPEN grant 875510], Janssen, Merck KGaA (aka EMD in Canada and US), Pfizer, Takeda and the Wellcome Trust [106169/ZZ14/Z].

